# Inferring the composition of a mixed culture of natural microbial isolates by deep sequencing

**DOI:** 10.1101/2024.08.05.606565

**Authors:** Mark Voorhies, Bastian Joehnk, Jessie Uehling, Keith Walcott, Claire Dubin, Heather L. Mead, Christina M. Homer, John N. Galgiani, Bridget M. Barker, Rachel B. Brem, Anita Sil

**Affiliations:** Department of Microbiology and Immunology, University of California San Francisco, San Francisco, California, United States of America; Department of Plant and Microbial Biology, University of California Berkeley, Berkeley, California, United States of America; The Pathogen and Microbiome Institute, Northern Arizona University, Flagstaff, AZ, USA; Division of Infectious Diseases, University of California San Francisco, San Francisco, California, United States of America; Valley Fever Center for Excellence, Department of Medicine, University of Arizona, Tucson, Arizona, United States of America; Chan Zuckerberg Biohub – San Francisco, San Francisco, California, United States of America

## Abstract

Next generation sequencing has unlocked a wealth of genotype information for microbial populations, but phenotyping remains a bottleneck for exploiting this information, particularly for pathogens that are difficult to manipulate. Here, we establish a method for high-throughput phenotyping of mixed cultures, in which the pattern of naturally occurring single-nucleotide polymorphisms in each isolate is used as intrinsic barcodes which can be read out by sequencing. We demonstrate that our method can correctly deconvolute strain proportions in simulated mixed-strain pools. As an experimental test of our method, we perform whole genome sequencing of 66 natural isolates of the thermally dimorphic pathogenic fungus *Coccidioides posadasii* and infer the strain compositions for large mixed pools of these strains after competition at 37°C and room temperature. We validate the results of these selection experiments by recapitulating the temperature-specific enrichment results in smaller pools. Additionally, we demonstrate that strain fitness estimated by our method can be used as a quantitative trait for genome-wide association studies. We anticipate that our method will be broadly applicable to natural populations of microbes and allow high-throughput phenotyping to match the rate of genomic data acquisition.

**Author summary:** The diversity of the gene pool in natural populations encodes a wealth of information about its molecular biology. This is an especially valuable resource for non-model organisms, from humans to many microbial pathogens, lacking traditional genetic approaches. An effective method for reading out this population genetic information is a genome wide association study (GWAS) which searches for genotypes correlated with a phenotype of interest. With the advent of cheap genotyping, high throughput phenotyping is the primary bottleneck for GWAS, particularly for microbes that are difficult to manipulate. Here, we take advantage of the fact that the naturally occurring genetic variation within each individual strain can be used as an intrinsic barcode, which can be used to read out relative abundance of each strain as a quantitative phenotype from a mixed culture. *Coccidioides posadasii*, the causative agent of Valley Fever, is a fungal pathogen that must be manipulated under biosafety level 3 conditions, precluding many high-throughput phenotyping approaches. We apply our method to pooled competitions of *C. posadasii* strains at environmental and host temperatures. We identify robustly growing and temperature-sensitive strains, confirm these inferences in validation pooled growth experiments, and successfully demonstrate their use in GWAS.

## Introduction

The rules by which genotype dictates phenotype are encoded in the genetic and phenotypic variation of natural populations. These rules can be decoded by statistical-genetic scans for polymorphisms that are co-inherited with, and potentially causal for, traits of interest among the progeny from controlled matings or among members of an outbred population. In many organismal systems, such efforts have been accelerated by pooled genotyping methods. This approach, originally called bulk segregant analysis in laboratory crosses [1] [2], has become an industry standard for invertebrate animals, eukaryotic microbes, and plants. The modern incarnation is to mix genetically distinct individuals, subject the resulting pool to selection for a phenotype of interest, and isolate DNA *en masse* from the subsets of the pool that pass or fail the selection [3]. From the resulting sequencing data, allele counts at each locus in turn then serve as input into statistical-genetic tests [4]. Against a backdrop of years of success, the pooled phenotyping-by-sequencing framework does have a key limitation: it does not quantify phenotypes of the individuals of the initial population. As a consequence, pooled methods preclude advanced statistical-genetic analyses at the haplotype and chromosomal level, including scans for genetic interactions between loci, calculations of polygenic risk scores, and population admixture control [5].

Strategies to infer individual strain abundances from sequencing of a complex pool have been developed in a related literature, that of microbial metagenomics. Can these tools be brought to bear on pooled statistical-genetic experiments? In metagenomics, a typical application requires simultaneous inference of each strain’s genotype and abundance in an ecological sample. To cut down on the resulting large search space, current methods make strong assumptions about pool membership (*e*.*g*. reference strains likely to be in the sample [6] [7], and/or small numbers of strains likely to dominate [8] [9] [10] [11]) whose validity in many cases may be unknown. But these caveats are not relevant in a statistical-genetic application using a pool of individuals whose genotype is known *a priori*. In such a scenario, inferring the prevalence of pool members from phenotyping-by-sequencing may be expected to work particularly well.

With this motivation, we set out to develop a method to fit the strain composition from full genome sequencing of a pooled sample, given individual genome sequencing of each member of that pool. As a test case for application of the approach, we focused on the fungal pathogen *Coccidioides posadasii. Coccidioides* species are the causative agents of Valley Fever and are endemic to California, Arizona, and other desert regions in the Americas [12]. *Coccidioides* grows as saprophytic hyphae in the environment [13]. The hyphae produce asexual spores, called arthroconidia, which, upon inhalation by mammalian hosts, convert to a unique pathogenic spherule morphology. The spherule undergoes internal segmentation to generate cells called endospores, which are released from spherules and presumably disseminate in the host. The genetic basis of these behaviors remains largely unknown. Many screening approaches that could fill the knowledge gap are out of reach because *Coccidioides* experiments can only be done under biosafety-level 3 containment laboratory conditions. Given the need for cheap and efficient experimental design in this system, forward genetics with phenotyping-by-sequencing methods is of particularly keen interest. We thus pioneered a scheme of pooling naturally varying strains of *C. posadasii* for growth and trait mapping. We implemented our model-fitting approach to infer strain abundance from the resulting data, and we then used relative abundances measured by this method as quantitative traits for association mapping, including control of population structure.

## Results

### Sequencing genetically distinct, clinical isolates of *Coccidioides posadasii*

With the ultimate goal of pooled growth assays and phenotyping-by-sequencing in *C. posadasii*, we sequenced 77 *C. posadasii* clinical isolates from Pima County, Arizona, of which 11 have been independently sequenced (Table S1), and we also resequenced the type strain of *C. posadasii*, Silveira [14]. The full set of sequencing data was combined with genomes of previously published strains to generate a phylogeny (Fig 1) which revealed that the Pima isolates in our pool corresponded to at least three previously identified populations [15].

**Fig 1.**
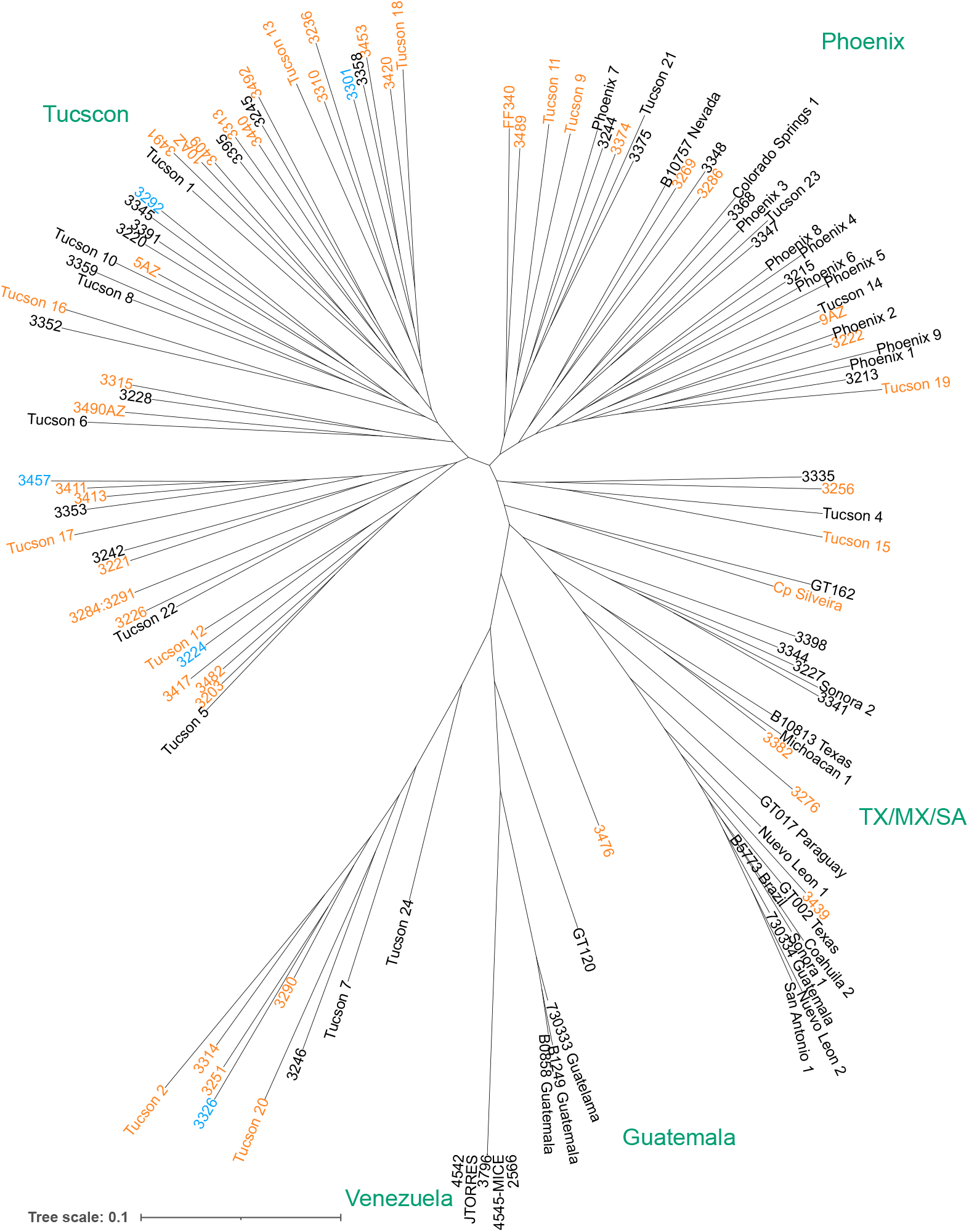
Phylogenetic distribution of *Coccidioides poasadasii* isolates. IQ-TREE inferred tree of the newly *C. posadasii* isolates relative to previously published strains. Strains included in the mixed culture experiments are highlighted in orange (55 strain pool only) or blue (55 strain and 5 strain pools).

As a model trait for pooled phenotyping assays, we chose a phenotype that could vary across our strains and be mapped by a genome-wide association study (GWAS) in a pooled format. Given that mammalian body temperature is both a cue for the fungal morphological transition and a stress that must be overcome to persist in the host, we chose differential growth at environmental (room temperature, RT) and host (37°C) temperatures as a useful test case for this purpose. As described in the following sections, we developed a method for fitting strain abundances to sequencing of a mixed culture seeded with a pool of *C. posadasii* strains, validated the method on simulated data, and we applied the method to real strain pools grown at 37°C or RT.

### Mixed pools can be deconvoluted by fitting a binomial model

For a pool of *M* strains that differ at *N* biallelic single-nucleotide polymorphisms (SNPs), we modeled the observed major allele read counts, *c*, out of total counts, *n*, at each variant position, *i*, based on a binomial distribution:

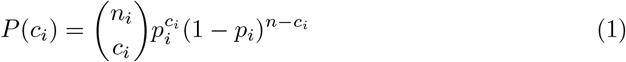

where the probability of observing the major allele, *p*_*i*_, is the total proportion of strains, *M*, harboring that allele:

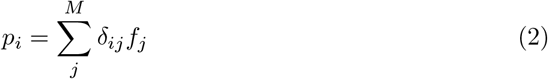

where *f*_*j*_ is the proportion of strain *j* and *δ*_*ij*_ = 1 for strains, *j*, with the major allele at position *i* and 0 otherwise.

The total likelihood of the observed read counts over all biallelic positions is then:

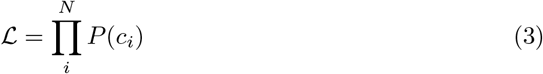

The unknown strain proportions can be estimated by finding the value of the vector, *F*, of strain proportions, *f*_*j*_, that maximizes the likelihood function. We found the maximum likelihood solution by minimizing a related objective function:

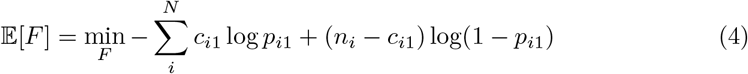

using simulated annealing [16, 17] (Fig S1).

An extended derivation of Eq 4 is given in the Materials and methods.

### Validation of strain abundance inference method by simulation

In order to validate our fitting method, we first simulated sequencing results from pooled growth of *C. posadasii* Silveira and 53 Pima County *C. posadasii* strains (the setup that we ultimately implemented experimentally; see below) by sampling reads from our single-isolate sequencing for chosen ground-truth proportions (Fig 2, top row). Given that we sampled from real sequencing runs, the simulations should incorporate the same positional and sequence biases, sequencing errors, etc., as in a real experiment. In particular, this simulation method is independent of the assumptions of our model and assumes only that there are no strain-specific biases in isolation of the DNA of the pool. We simulated mock pools at a depth of 30 million reads, a lower limit of the sequencing depth of our true pooled experiment below.

**Fig 2.**
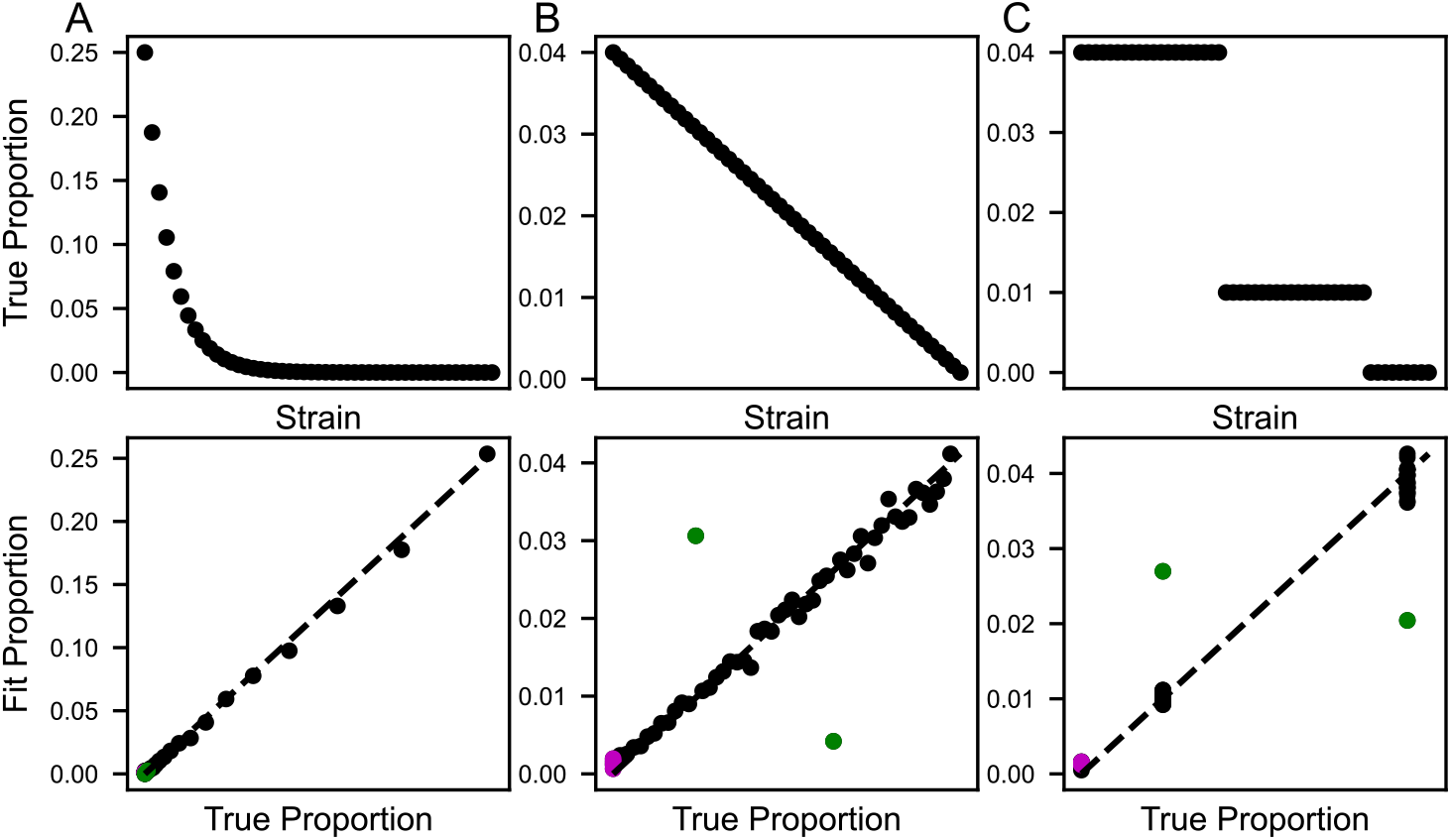
Validation of model with simulated data. Shown are results from simulations of three pools with different strain proportions (A, B, and C). In the top row of plots, each point reports the true abundance of one strain in the respective simulated pool, as a proportion of total biomass. In the bottom row of plots, each point reports the abundance of one strain fit by the model (y-axis) and the true simulated abundance (x-axis) for the respective simulated pool. Clonal pair (strains 3284 and 3291) indicated in green. Strains absent from the simulated pools indicated in magenta.

2800 steps of simulated annealing were sufficient for the fit to converge in each case (Fig S2), and the fit strain proportions recapitulated ground truth to within 1% (Fig 2, bottom row). As expected, the exception to the rule was a pair of genetically indistinguishable clonal strains present in the true strain set (green circles in Fig 2B and see Methods). Similar results were obtained when we used only a transposon-free region between the *KU70* and *HSF1* genes (coordinates: CP075070.1:4382963..5065814), representing about 5% of the genome (Fig S2).

### Inferring strain abundances from experimental pooled sequencing

To apply our approach to real data, we performed a first set of competition experiments for 54 well-germinating strains of *C. posadasii* at host (37°C) and environmental (RT) temperatures.

The procedure was repeated in two batches, with three pools grown at each temperature for each batch, for a total of 12 pools. In each case we inoculated arthroconidia into liquid culture and incubated to allow germination and hyphal growth. We then isolated DNA from each, carried out sequencing, quantified alleles at SNPs, and inferred strain abundance with our fitting method. In each fit, simulated annealing converged (Fig S3). Inferred strain abundances varied more among the pools cultured at 37°C than those at RT (Fig 3A), but significance testing was still highly powered to resolve temperature differences in abundance for eight strains (blue and red points, Fig 3B). Two other strains exhibited evidence for jackpotting in the cultures, with very high abundance independent of temperature (purple points, Fig 3B).

**Fig 3.**
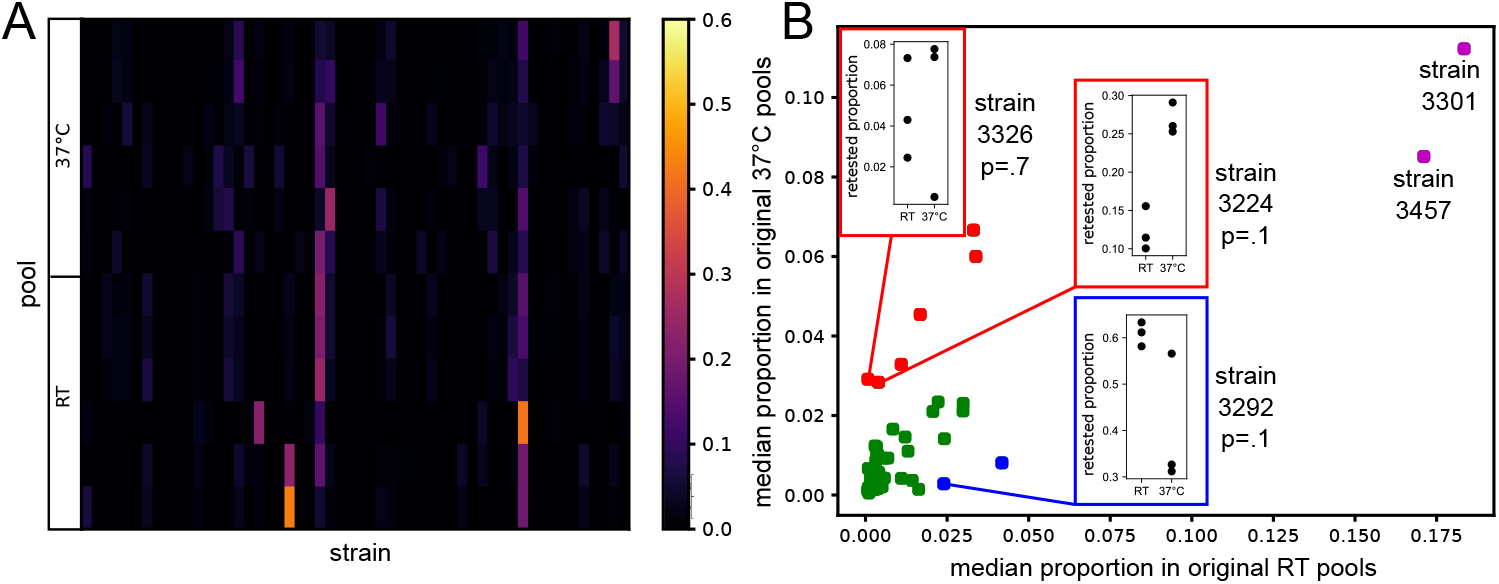
Strain proportions fit from real pool sequencing. (A) Heatmap showing fit proportions for each of 54 strains (columns) in each of 12 pooled liquid cultures (rows) grown for 14 days at 37°C or RT. (B) In the main plot, each point reports abundance as a median across replicate pools of the indicated temperatures from (A). Strains are colored based on whether they were: highly abundant at both temperatures (purple), enriched at 37°C (red), enriched at RT (blue), or not detectably temperature-dependent (green). Insets show proportions for strains 3326, 3224 and 3292 when retested in a 5 strain pool (p-values from Wilcoxon tests).

In order to validate the strain-specific temperature biases measured by our inference approach, we carried out a second, smaller pooled competition experiment consisting of the two *C. posadasii* strains that had dominated cultures in the first round (3301 and 3457), two of the 37°C enriched strains from the first round (3224 and 3326), and one of the RT enriched strains from the first round (3292). We grew the smaller pools in triplicate at 37°C and RT and again isolated and sequenced DNA from each replicate. Running our inference method on the resulting sequencing data correctly identified the five strains present in these pools (Fig S4). Furthermore, the temperature-dependent abundance patterns in this experiment recapitulated the trends we had seen in the first round, with strains 3224 and 3292 again exhibiting 37°C and RT biases, respectively (insets, Fig 3B). As in the larger pools, strains 3301 and 3457 did not show a temperature bias in abundance. They did not, however, show evidence for jackpotting in the smaller pools. Instead, in the latter, strain 3292 was the most abundant strain at either temperature (insets, Fig 3B; Fig S4). Taken together, these results show that despite variation in jackpotting effects across experimental designs, strain differences in growth rate between conditions are reproducible across different *C. posadasii* pool compositions and can be inferred robustly with our fitting approach. In principle, strain variation in temperature tolerance could derive from differences in the ability to cope with thermal stress or to differential response to the cue of host temperature.

### Application to GWAS

We reasoned that the inferred abundances from our larger pooled growth experiment could be used as the basis for genetic dissection of variation in temperature-dependent growth, via GWAS. Given the population structure in our sampled *C. posadasii* strain set (Fig 1), our association test used a linear mixed model as implemented in GEMMA [18] to correct for kinship effects. We formulated the phenotype of each strain as the log fold-change of the difference in inferred abundance between pooled growth experiments at RT and 37°C. A genomic scan for variants associated with this trait revealed a single locus with 12 associated SNPs passing a 10% false discovery rate (7 SNPs at nominal p = 2.18e-05 and adjusted p 0.083; 5 SNPs at nominal p = 2.5e-05 and adjusted p = 0.1; Fig 4A-C). The same locus emerged as the top hit from association test schemes that did not correct for population structure, including a linear model (LM) with no kinship terms (nominal p = 2.18e-05) and a non-parametric Wilcoxon test (nominal p = 9.69e-6), attesting to the robustness of the signal irrespective of admixture effects. All of the associated SNPs at the locus fell in a single predicted gene of unknown function, D8B26_001557, unique to the Onygenales. In expression profiles of the *C. posadasii* type strain [19], this gene was induced in the pathogenic spherule form of the fungus relative to the vegetative hyphal form and depended on the transcription factor Ryp1, a driver of temperature-evoked development in *C. posadasii* [19] and other fungal pathogens (Fig 4F).

**Fig 4.**
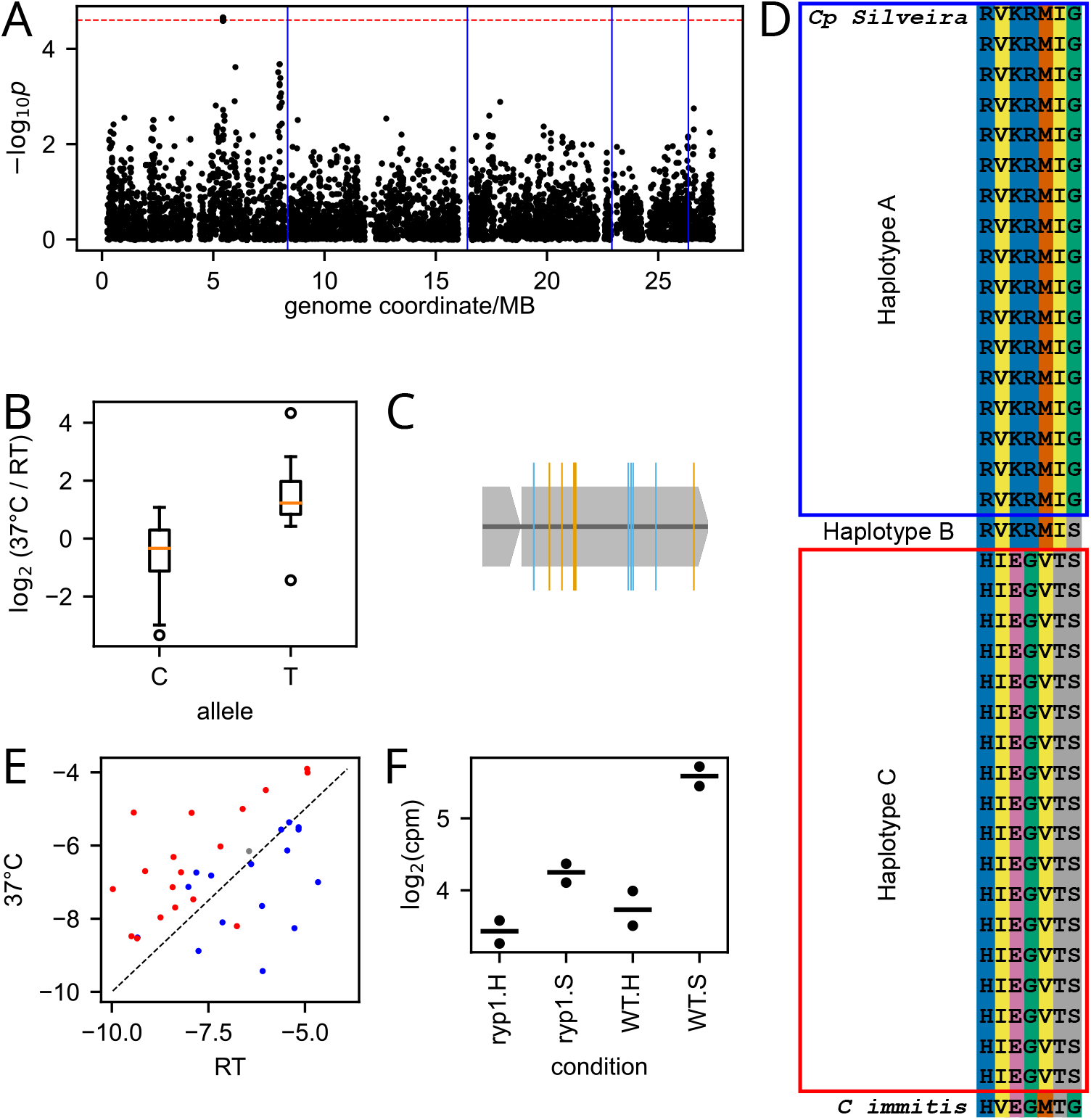
Genome-wide assocation study (GWAS) of temperature-dependent growth inferrred from phenotyping-by-sequencing. (A) Each point reports results from a test of association of one single-nucleotide polymorphism (SNP) with inferred temperature-dependent growth across *C. posadasii* strains. The y-axis reports the uncorrected p-value from an assocation test via a linear mixed model in the GEMMA package with admixture correction, and the x-axis reports genomic location. Horizontal dashed red line indicates 10% false discovery rate based on 1000 permutations of the phenotype vector. Blue vertical lines indicate chromosome boundaries. (B) Each column reports the distribution of inferred temperature-dependent phenotypes among strains harboring the indicated allele at the top GWAS hit locus, in D8B26_001557. (C) Schematic of D8B26_001557, with locations of SNPs whose association significance passed a 10% false discovery rate threshold in orange (missense) or blue (silent). (D) Each row reports the sequence in one *C. posadasii* strain (or *C. immitis*, at bottom) at the seven non-silent GWAS hit sites from (C). Blank labels at left indicate strains genotyped in this study. (E) Each point reports the log of the median of the inferred abundance for one strain at the indicated temperature, colored by its inheritance at the top hit in D8B26_001557 as in (D). (F) Transcript levels of D8B26_001557 in WT or *ryp1* mutant in spherule-inducing (S) or hyphal-inducing (H) conditions (data from [19]).

In our association results, seven of the SNPs in D8B26_001557 associated with temperature-dependent strain abundance drove non-synonymous changes (Fig 4D). They defined three haplotypes across our Pima County *C. posadasii* strain set (Fig 4D and 4E): haplotype A, associated with higher abundance at RT and present in 16 of the strains; haplotype C, associated with higher abundance at 37°C and present in 18 strains ; and haplotype B, generated by a single crossover between haplotypes A and C and present in a single strain with similar abundance at both temperatures. Comparison against four strains of the pathogen relative *C. immitis* (RS, RMSCC 2394, RMSCC 3703, and H538.4) revealed an invariant haplotype in *C. immitis*, distinct from the *C. posadasii* haplotypes (Fig 4D).

We conclude that variants in D8B26_001557 represent a compelling candidate determinant of the variation in temperature-dependent growth across our *C. posadasii* strains. This discovery validates our pipeline of pooled growth, phenotyping-by-sequencing in pools, and inferred strain abundance as a highly-powered strategy for statistical genetics.

## Discussion

Pooling approaches for statistical genetics, though powerful, typically preclude the use of chromosome- or haplotype-level advanced mapping methods. Here, we adapt mathematical techniques originally developed for deconvolution of strain data from metagenomic sequencing to infer the relative proportions of strains of a single species in a mixed culture. We show that this approach can be used to score quantitative phenotypes for differential growth between two conditions in a pooled format and that these phenotypes can be applied to GWAS in *Coccidioides*.

Progress in the molecular understanding of microbial pathogens is often hampered by the difficulty of genetic screening tools. Indeed, our work represents a first-ever whole-genome functional-genomic screen in *Coccidioides*. To date, only 10 genes related to virulence [20] [21] [22] [23] [24] [25] [26] [27] and morphology [19] [28] are known in this organism, most initially identified via secretion from the pathogenic spherule form [20] [21] [22] [23] [24] [25] [26] [27] or by hypotheses based on the biology of other fungi [24] [25] [26] [27] [19] [28]. Forward genetic screens are likely to be of great utility in advancing the field, and a recent small-scale screen of 24 *Coccidioides* insertional mutants for virulence in *Galleria* serves as an additional foundation for this principle [29]. Given the success of GWAS in fungal model systems, plant pathogens and commensals, and opportunistic animal pathogens, we expect the natural variation-based approach to be equally powerful in human pathogens, especially with the pooled growth paradigm we establish here.

As temperature is both an important developmental cue and a stress inherent to the host environment, we chose growth of *Coccidioides* at 37°C as an easily controlled model trait to test our pooled GWAS method. Our inference of variation in temperature-dependent growth across *C. posadasii* strains from the pooled experimental format is consistent with a previous survey of growth across temperatures in a strain-by-strain setting [30], and our discovery of D8B26_001557 as a candidate determinant of these differences serves as an additional proof of concept for our approach. Regulation of this gene by the Ryp1 transcription factor is consistent with the control of its ortholog in *Histoplasma ohiense* G217B, I7I48_06129^1^, by *Histoplasma* Ryp1 [31]. Ryp1 is a master regulator of the temperature-dependent transition of *Histoplasma* from hyphae to yeast [32] [19] and is likewise required for spherule formation in *Coccidioides* [19]. Given the role of Ryp1 in temperature-dependent transcriptional regulation, it is appealing to discover a Ryp1 target associated with temperature-dependent growth. Of additional interest is that fact that the Ryp1 regulon is associated with pathogenesis in many fungi [33] [34] [35] [36] [37] [38] [39] [40]. Many of the known virulence factors of *Histoplasma* are direct Ryp1 targets [31], and the *Coccidioides* virulence factors SOWgp, MEP1, and urease are Ryp1 regulated [19]. Thus it is tempting to speculate that our GWAS hit locus, D8B26_001557, may ultimately prove to have relevance for virulence behaviors. We observe that *C. posadasii* Silveira has the haplotype that favors growth at RT, consistent with the previous observation of compromised growth at high temperature for this strain [30].

Our GWAS of temperature-dependent growth in *C. posadasii* from pooled experimental measurements opens a window onto the study of natural variation in other virulence-relevant traits in this pathogen, including development of infectious spherules. More generally, our strategy should be applicable to phenotyping of pools of genetically distinct strain sets in many organismal systems. Our current likelihood function assumes a haploid genome but could readily be extended to the diploid case. The approach is expected to have particular impact in statistical-genetic scans that use population structure correction, as we have implemented it here for *C. posadasii*, and other multi-locus mapping tools. Our method will also be of use in applications beyond statistical genetics *per se* – for example, correlation analysis of multiple phenotypes or rapid surveys of a set of isolates to select strains with extreme phenotypes for comparative ‘omics.

In summary, the pooled phenotyping format has become a linchpin of the field, especially for non-model systems like pathogens; and with adaptations like ours, powerful genomic methods, even those originally developed for classical phenotyping on one individual at a time, come within reach for the cheap, expedient pooled experimental design.

## Materials and methods

### Strains and growth conditions

All strains were received from J.G. and collected from a single hospital site in Tucson (Southern Arizona Veterans Administration Health Care Service).

Strains were transferred from the B.M.B. lab at NAU to the A.S. lab at UCSF as follows: Slants containing 2xGYE (2% glucose, 1% yeast extract) were inoculated with glycerol stocks (1% glucose, 1% yeast extract, 10% glycerol) of previously harvested arthroconidia.

Arthroconidia from 54 isolates of *C. posadasii* used in this study were generated by cultivation on solid glucose yeast extract medium (GYE 2X: glucose 20 g/L, yeast extract 10 g/L, agar 15 g/L) in 125 ml vented suspension flasks for 4-6 weeks at 30°C. Arthroconidia were harvested using phosphate buffer (PBS) and a cell scraper to dislodge arthroconidia from grown mycelium. The hyphal/spore mixture was filtered through miracloth to isolate the arthroconidia from hyphal mass. The spores were washed twice with PBS before being stored at 4°C for long term storage. The spore solutions were quantified with a hemocytometer and adjusted to working concentrations of 105 arthroconidia/µL. An additional isolate, *C. posadasii* RMSCC 1043, did not germinate well and did not yield sufficient material for sequencing. We eliminated RMSCC 1043 from simulation studies and analyses of real experimental data (see below) under the assumption that it would not contribute to mycelial pools.

### Pooled competition experiments

Arthroconidia from *C. posadasii* Silveira and 53 clinical isolates of *C. posadasii* (plus RMSCC 1043) were grown in competition under conditions associated with the host or the environment. For each clinical isolate, 12.74 µL of the 105 arthroconidia/µL stocks was added to 6.3 mL of PBS in a 15mL conical, resulting in a total spore concentration of 107 spores/ml. Six flasks containing 25mL of GYE 2X liquid media were inoculated with 550 µL of the spore mixture, representing 105 spores per isolate in each culture. The cultures were incubated for 14 days on an orbital shaker at 120 RPM either at room temperature or at 37°C in the presence of 5% CO2.

The mycelium from each culture was collected in miracloth filters, washed with PBS to remove carryover media, then pat dried on paper towel to remove excess moisture. Dried mycelium from each culture was subsequently frozen in liquid nitrogen and pulverized by cryo milling with a Retsch MM400 (30 Hz for 1 minute). 0.02-0.06 g pulverized frozen mycelium was transferred to a 2 mL screw cap tube containing 700 µL lysis buffer (0.05 M Tris pH 7.2, 0.05 M EDTA, 3% SDS, 1% *β*-mercaptoethanol) thoroughly mixed and heated for 1 hour at 65°C. Subsequently 800 µL phenol:isopropanol:isoamylalcohol (25:24:1) was added to each tube and mixed by inverting several times. Tubes were centrifuged at (max speed) for 15 minutes and genomic DNA was precipitated from the aqueous phase by pipetting it into and mixing with 450 µl 2-propanol + 20 µl sodium acetate (3 M) then pelleted by centrifugation at (max speed) for two minutes. The DNA pellets were washed twice with 70% ethanol then dried at 50°C for 5-10 minutes. DNA was eluted into 200 µl TE 1X (10 mM Tris, 1 mM EDTA, pH 8) + 2 µl RNAse A (10 mg/mL) then stored at -20°C.

### NGS-library preparation and sequencing

Genomic DNA from each sample was prepared for next generation sequencing using the Nextera DNA Flex Library Prep kit (Illumina). In brief, 500 ng of gDNA from each pooled competition experiment were used as input and tagmented (process to fragment and tag the DNA with adapter sequences) with Bead-Linked Transposomes. After washing, the tagmented DNA was amplified with unique combinations of i7 and i5 index adapters. Library qualities were assessed using an Agilent High Sensitivity DNA chip run on a 2100 Bioanalyzer Instrument. Equal molar amounts of each library were multiplexed to a final concentration of 5-10 nM.

Individual strains were multiplexed in four batches and sequenced on the Illumina HiSeq 4000 platform at the UCB QB3 core or at the UCSF Center for Advanced Technology (CAT).

Pooled cultures were multiplexed in three batches and sequenced on the Illumina HiSeq 4000 platform at the UCSF CAT or the Illumina NovaSeq S2 platform at the Chan Zuckerberg Biohub.

### Simulations

Mock pools were simulated by sampling read pairs from the individual isolates in the required proportions by reservoir sampling (“Algorithm R” of Vitter [41]).

### Phylogenetics

FASTQ files for previously published *C. posadasii* strains [42] [15] were downloaded from SRA:PRJNA274372 and SRA:PRJNA438145. The *C. posadasii* reference genome and annotations [43] were downloaded from BioProject PRJNA664774. We used RepeatMasker 4.1.2 [44] to identify regions of the *C. posadasii* reference genome with low complexity or simple repeats, or mapping to a library of known *Coccidioides* transposable elements [45].

For each isolate and GWAS pool, we aligned reads from the FASTQ file to the *C. posadasii* reference genome using BWA MEM 0.7.17 [46], then sorted and indexed the aligned reads using SAMTOOLS 1.8 [47]. We used PILON 1.23 [48] with the –variant flag to create VCF (Variant Call Format [49]) files from the aligned reads. Variant sites in each VCF were filtered as follows: variants annotated as “LowCoverage” by pilon or with depth greater than three times the average depth of the sample were removed. We also removed variants within the repetitive regions identified by RepeatMasker.

Vcf2phylip.py (https://doi.org/10.5281/zenodo.2540861) was used to convert from a VCF containing all biallelic sites with minor allele frequency of at least 5% and no missing data to phylip format. We used this file as input to the ModelFinderPlus model of IQ-TREE 1.6.12 [50] [51] [52], which identified the TVM+F+ASC+R5 model as optimal using Bayesian information criterion. We bootstrapped this model with 1000 replicates using the SH-like approximate likelihood ratio test and ultrafast bootstrap approximation methods. The consensus tree from the ultrafast bootstrap approximation was visualized with iTOL [53]. The 12 resequenced strains were included in the tree inference, were correctly identified as redundant by IQ-TREE, and are not shown in Fig 1.

### Read alignment

Paired-end reads from individual isolates and simulated and real pools were aligned to the *Coccidioides posadasii* var. Silveira genome assembly using BWA MEM [46] with default parameters.

### SNP calling from individual isolates

For the purpose of fitting proportions in simulated and real pools, SNPs were defined from the individual isolate read alignments as nucleotide positions covered by at least 10 reads from each isolate, with at least 85% of the reads from each isolate supporting a single allele, and with exactly two total alleles over all isolates.

### Model and fitting procedure

We give here an extended derivation of the objective function from Eq 4 from the Results section.

Given major allele counts, *c*_*i*_, for each biallelic position, *i*, out of *N* total biallelic positions over *M* strains, we would like to find the strain frequencies *F* that best fit the allele counts under the constraints that the frequencies are positive, 0 *≤ f*_*j*_ *≤* 1*∀j*, and sum to 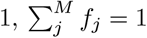.

The probability that a count at *i* is due to strain *j* is its frequency:

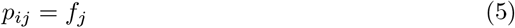

Therefore, the probability of observing *c*_*i*_ counts at a position where strain *j* is the *only* strain with the major allele is given by the binomial distribution:

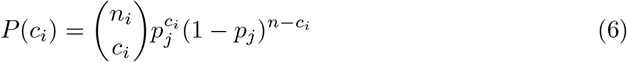

For positions where more than one strain has the major allele we need to account for all of the different ways that the *c*_*i*_ counts could be partitioned among the strains. As this is already built into Eq 6 via the binomial coefficient, it is easiest to first sum the individual strain probabilities and then insert this total major allele frequency into that equation:

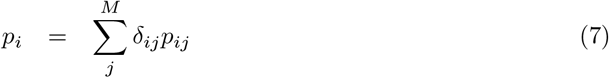

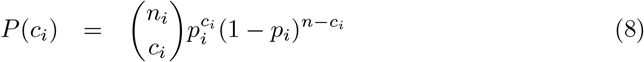

where *δ*_*ij*_ is 1 if strain *j* has the major allele at *i* and 0 otherwise.

The total likelihood of the observations, *C*, given a candidate solution, *F*, is:

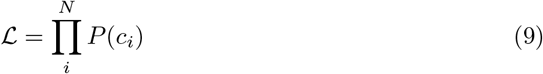

which can be maximized by minimizing the negative log likelihood:

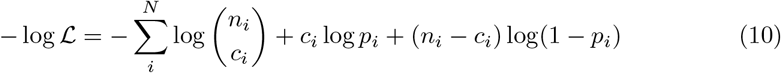

Note that the binomial coefficient is a fixed term that depends only on the observed allele ratio, so it can be dropped for the minimization:

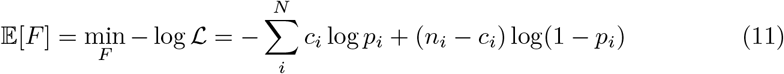

which is Eq 4 from the Results section.

### GWAS

For GWAS, we considered all strains in the pool except for RMSCC 1043 (not sequenced), *C. posadasii* (not a Pima isolate), and the two jackpotting strains (3301 and 3457), leaving 51 strains. We selected a total of 8552 SNPs outside of LTR transposon-rich regions (defined as in [54]) and in coding sequence with at least 25% minor allele frequency (MAF) relative to the 51 analyzed strains.

Per strain phenotypes were calculated as the difference in median log_2_ proportions between 37°C and RT.

GWAS analysis was then carried out using GEMMA [18] 0.98.4 to infer the kinship matrix from the SNP matrix:

~~~
gemma -g genotypes.bimbam.gz -p phenotypes.txt -gk \
 -outdir gwas -o kinship
~~~

to fit a full linear mixed model (LMM):

~~~
gemma -g genotypes.bimbam.gz -p phenotypes.txt -n 1 \
 -k gwas/kinship.cXX.txt -lmm 4 -outdir gwas -o lmm4
~~~

and to fit an equivalent linear model (LM) without random effects *(i*.*e*., *assuming no population structure)*:

~~~
gemma -g genotypes.bimbam.gz -p phenotypes.txt -n 1 \
 -k gwas/kinship.cXX.txt -lm 4 -outdir gwas -o lm4
~~~

We likewise carried out GWAS assuming no population structure and without the assumption of linearity by using a Wilcoxon test as implemented in R [55].

p-values from the LMM fit were corrected for multiple hypothesis testing by re-running the analysis for 1000 random permutations of the phenotype vector and counting the frequency at which the unadjusted p-values occurred in these permuted controls.

Protein sequences for the *C. immitis* orthologs of D8B26_001557 were obtained from GenBank (GCA_000149895.1_ASM14989v1 RMSCC 2394, GCA_000150085.1_ASM15008v1 RMSCC 3703, GCF_000149335.2_ASM14933v2 RS, and GCA_000149815.1_ASM14981v1 H538.4) and aligned to the *C. posadasii* sequence with PROBCONS 1.12 [56].

**Fig S1.**
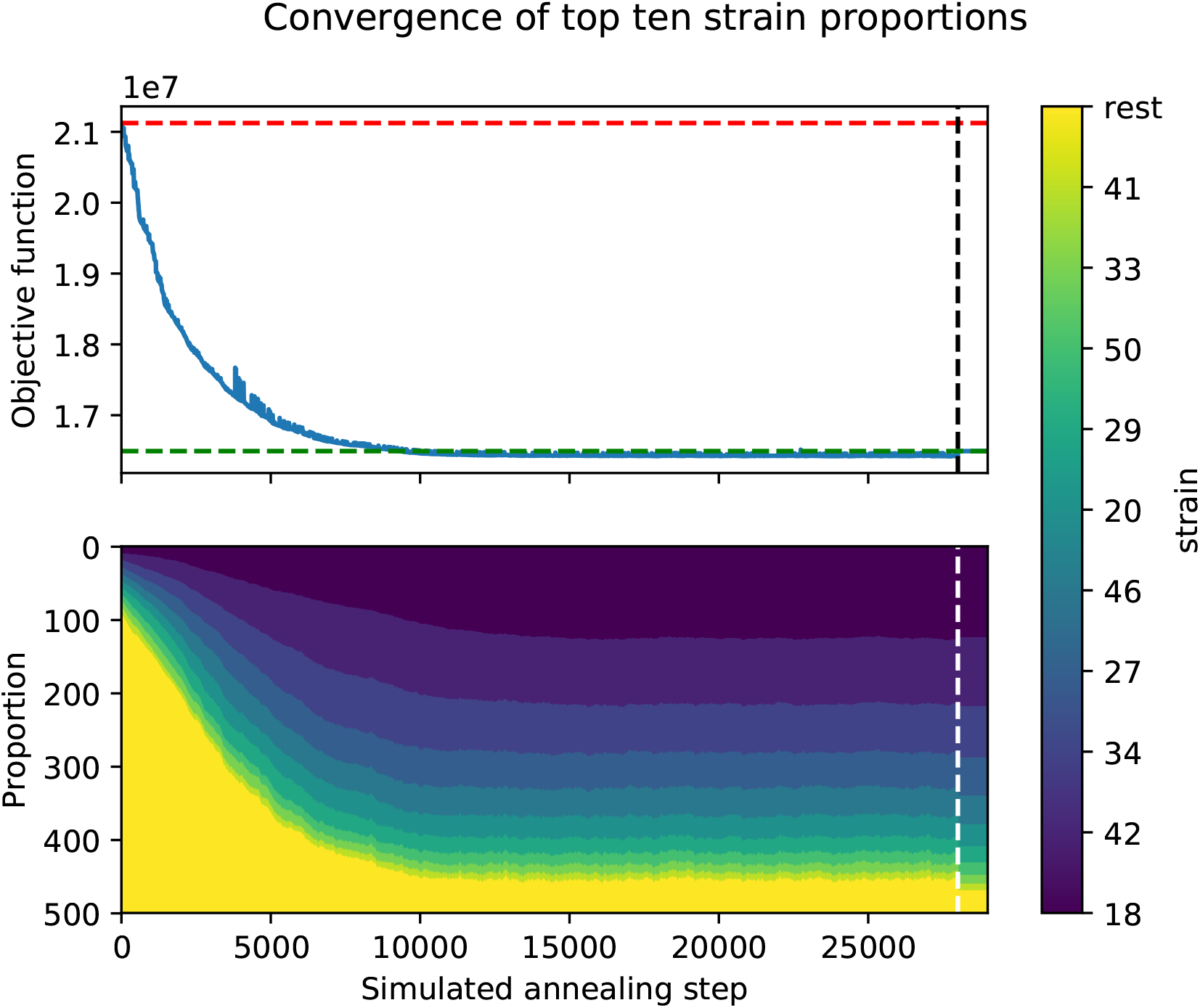
Fitting strain proportions with simulated annealing. (Top) Objective function for each simulated annealing step for a simulated pool (blue line). Values of the objective function are shown as dashed lines for a uniform strain distribution (red) and for the known correct distribution (green). (Bottom) Estimated proportions of the top ten strains or the remaining strains (“rest”) plotted as stacked relative proportions for each simulated annealing step. For both the top and bottom plots, the true solution is plotted to the right of the vertial dashed line.

## Supporting information

**S1 Table**. Table of strains used in this work as tab-delimited text file. Publication_ID gives the strain name used in the main text and figures. Additional identifiers that have been associated with a strain are given in Preliminary_ID, Collection_ID, ALT_ID, UCSF_ID, and FASTQ_prefix. Collection details are given in SPECIES, COUNTRY, CITY/LOCATION, STATE/COUNTRY, ISOLATION/DISEASE_INFO, and YEAR. Strains used in the first set of pools, the retesting set of pool, or the GWAS analysis are indicated with “True” in the Pool1, Pool2, or GWAS columns respectively. Previously sequenced strains are indicated by SRA run ID in the Previously_sequenced column. Strains newly sequenced or resequenced in this work are indicated with “True” in the Sequenced column.

**Fig S2.**
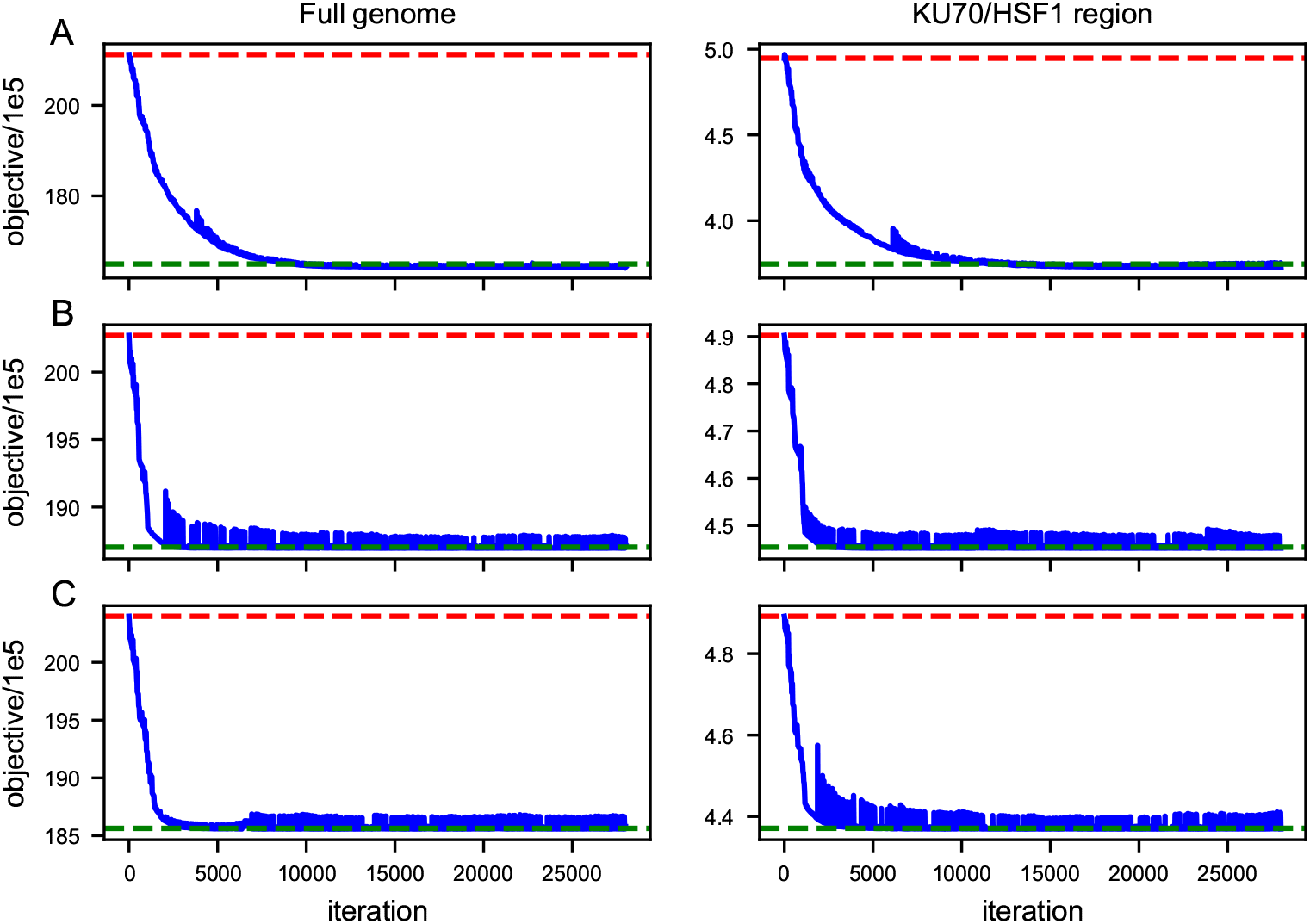
Simulated annealing converges to ground truth for simulated pools. Objective function at each simulated annealing step for the simulated pools A, B, and C (blue). Values of the objective function are shown as dashed lines for a uniform strain distribution (red) and for the known correct distribution (green). Curves are plotted for SNPs identified from the full genome (left) or from the transposon-free region between KU70 and HSF1, representing about 5% of the genome.

**S2 File. Code**. Zip archive of the python code implementing our fitting method and the Jupyter notebooks required to generate the figures in the paper.

**Fig S3.**
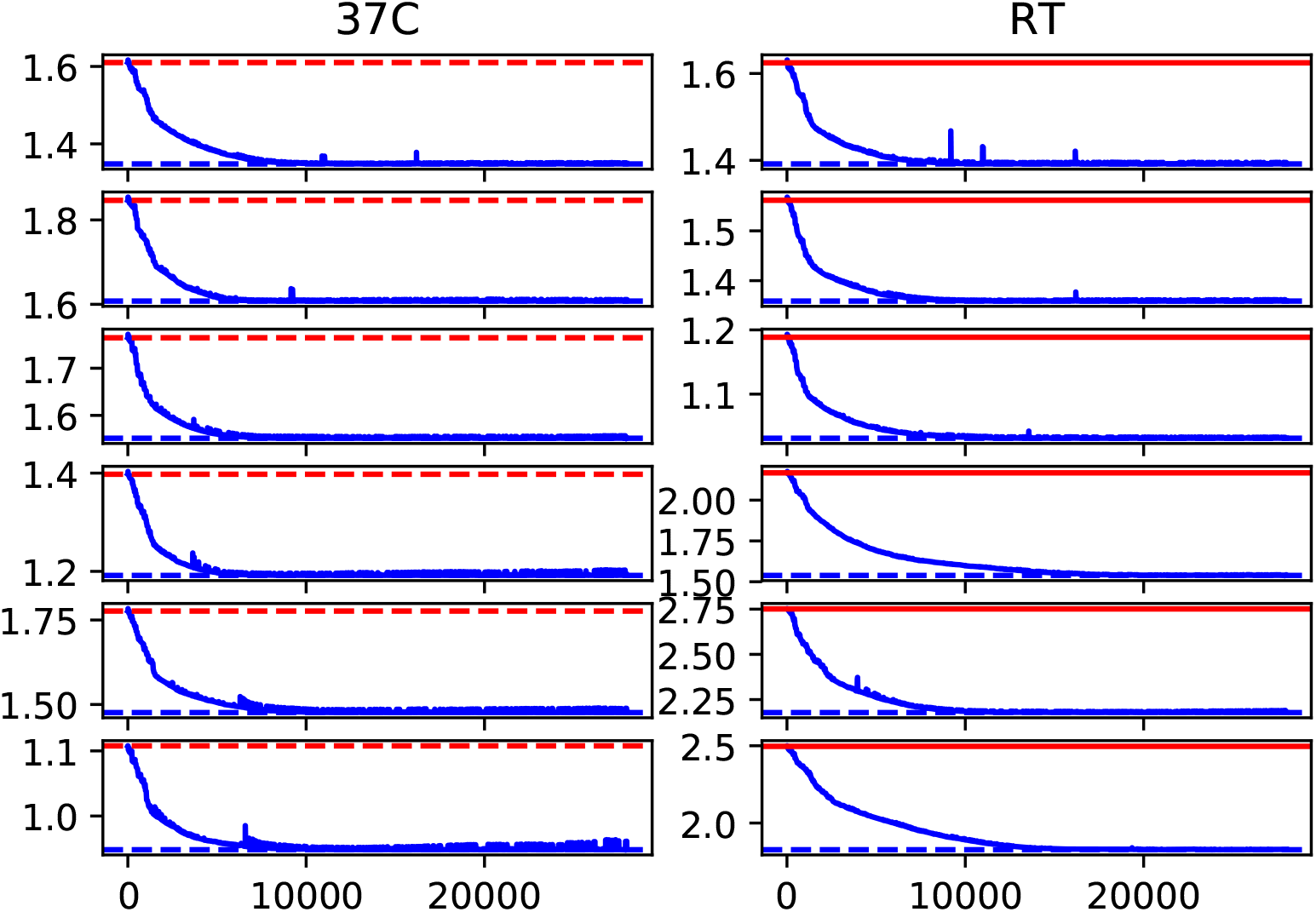
Simulated annealing converges for real pools. For each of 12 pools, the objective function is plotted as a function of the simulated annealing step as a solid blue line with the minimum value as a dashed blue line and the objective function evaluated for a uniform strain distribution plotted as a solid red line.

**Fig S4.**
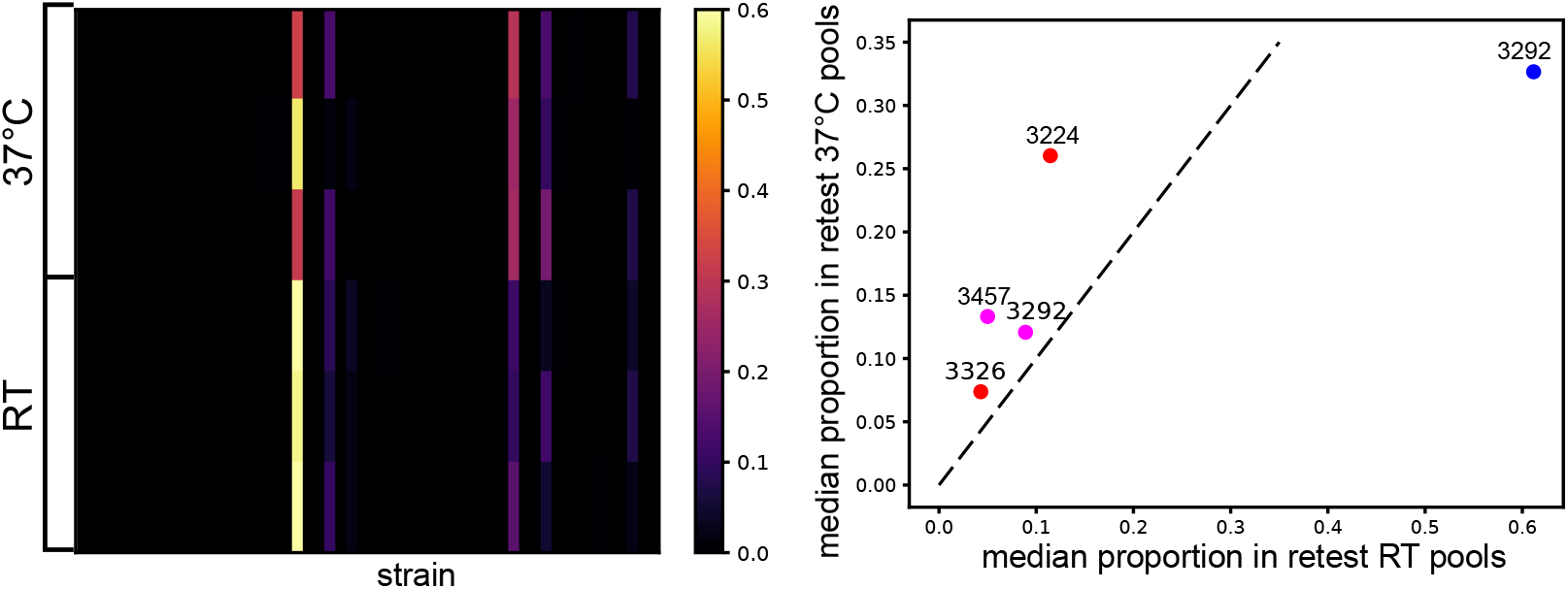
Strain proportions fit from retest pool sequencing. (A) Heatmap showing fit proportions for each of 54 strains (columns) in each of 6 pooled liquid cultures of 5 strains (rows) grown for 14 days at 37°C or RT. (B) Scatter plot of median proportion for each strain in the 37°C vs. RT pools from (A). Strains are colored as in Fig 3B. Dashed line indicates a slope of 1.

## Data availability

Sequencing reads have been deposited in the NCBI short read archive (SRA) under accessions PRJNA1143091 (individual strains) and PRJNA1143168 (pools).

Remaining data (SNP matrices and simulated pools) will be deposited in Data Dryad upon publication.

## Acknowledgments

We thank the UCSF CAT, UCB QB3, and Chan Zuckerberg Biohub for DNA sequencing. We thank members of the Sil, Brem, Noble, and Madhani labs for useful discussions.

This research was supported by the HHMI Hanna Gray Fellowship (to CH), the Program for Breakthrough Biomedical Research, which is partially funded by the Sandler Foundation (to CH), NIH R21AI172185 (to AS), and NIH U19AI166798 (to AS, RB, and BB). The UCSF CAT is supported by UCSF PBBR, RRP IMIA, and NIH 1S10OD028511-01 grants.

## Footnotes

1 HISTO_ZT.Contig1089.Fgenesh_histo.56.final_new in [31]

## References

1. Giovannoni J, Wing R, Ganal M, Tanksley S. Isolation of molecular markers from specific chromosomal intervals using DNA pools from existing mapping populations. Nucleic acids research. 1991;19:6553–8.

2. Michelmore R, Paran I, Kesseli R. Identification of markers linked to disease-resistance genes by bulked segregant analysis: a rapid method to detect markers in specific genomic regions by using segregating populations. Proceedings of the National Academy of Sciences of the United States of America. 1991;88:9828–32.

3. Macgregor S, Zhao Z, Henders A, Nicholas M, Montgomery G, Visscher P. Highly cost-efficient genome-wide association studies using DNA pools and dense SNP arrays. Nucleic acids research. 2019;36(6):e35.

4. Zou C, Wang P, Xu Y. Bulked sample analysis in genetics, genomics and crop improvement. Plant biotechnology journal. 2016;14:1941–55.

5. Brandes N, Weissbrod O, Linial M. Open problems in human trait genetics. Genome biology. 2022;23:131.

6. Albanese D, Donati C. Strain profiling and epidemiology of bacterial species from metagenomic sequencing. Nature communications. 2018;8(1):2260.

7. Ahn T, Chai J, Pan C. Sigma: strain-level inference of genomes from metagenomic analysis for biosurveillance. Bioinformatics (Oxford, England). 2021;31(2):170–7.

8. Smith B, Li X, Shi Z, Abate A, Pollard K. Scalable Microbial Strain Inference in Metagenomic Data Using StrainFacts. Frontiers in bioinformatics. 2022;2:867386.

9. Smillie C, Sauk J, Gevers D, Friedman J, Sung J, Youngster I, et al. Strain Tracking Reveals the Determinants of Bacterial Engraftment in the Human Gut Following Fecal Microbiota Transplantation. Cell host & microbe. 2022;23(2):229–240.e5.

10. O’Brien J, Didelot X, Iqbal Z, Amenga-Etego L, Ahiska B, Falush D. A Bayesian approach to inferring the phylogenetic structure of communities from metagenomic data. Genetics. 2022;197(3):925–37.

11. Luo C, Knight R, Siljander H, Knip M, Xavier R, Gevers D. ConStrains identifies microbial strains in metagenomic datasets. Nature biotechnology. 2024;33(10):1045–52.

12. Johnson L, Gaab E, Sanchez J, Bui P, Nobile C, Hoyer K, et al. Valley fever: danger lurking in a dust cloud. Microbes and infection. 2014;16:591–600.

13. Cole G, Sun S. Arthroconidium-Spherule-Endospore Transformation in Coccidioides immitis. In: Szaniszlo P, Harris J, editors. Fungal Dimorphism: With Emphasis on Fungi Pathogenic for Humans. Boston, MA: Springer US; 1985. p. 281–333. Available from: 10.1007/978-1-4684-4982-2_12.

14. Fisher M, Koenig G, White T, Taylor J. Molecular and phenotypic description of Coccidioides posadasii sp. nov., previously recognized as the non-California population of Coccidioides immitis. Mycologia. 2002;94(1):73–84.

15. Teixeira M, Alvarado P, Roe C, 3rd GT, Patane J, Sahl J, et al. Population Structure and Genetic Diversity among Isolates of Coccidioides posadasii in Venezuela and Surrounding Regions. mBio. 2019;10(6):emBio.01976–19.

16. Metropolis N, Rosenbluth A, Rosenbluth M, Teller A, Teller E. Equations of state calculations by fast computing machines. J Chem Phys. 1953;21:1087 – 1092.

17. Press WH, Teukolsky SA, Vetterling WT, Flannery BP. Numerical Recipes in C, 2nd ed. Press Syndicate of the University of Cambridge; 1992. Available from: http://www.nr.com/ http://lib-www.lanl.gov/numerical/bookcpdf.html.

18. Zhou X, Stephens M. Genome-wide efficient mixed-model analysis for association studies. Nature genetics. 2012;44(7):821–4.

19. Mandel M, Beyhan S, Voorhies M, Shubitz L, Galgiani J, Orbach M, et al. The WOPR family protein Ryp1 is a key regulator of gene expression, development, and virulence in the thermally dimorphic fungal pathogen Coccidioides posadasii. PLoS pathogens. 2022;18(4):e1009832.

20. Hung C, Yu J, Seshan K, Reichard U, Cole G. A parasitic phase-specific adhesin of Coccidioides immitis contributes to the virulence of this respiratory Fungal pathogen. Infection and immunity. 2002;70:3443–56.

21. Hung C, Seshan K, Yu J, Schaller R, Xue J, Basrur V, et al. A metalloproteinase of Coccidioides posadasii contributes to evasion of host detection. Infection and immunity. 2005;73:6689–703.

22. Mirbod-Donovan F, Schaller R, Hung C, Xue J, Reichard U, Cole G. Urease produced by Coccidioides posadasii contributes to the virulence of this respiratory pathogen. Infection and immunity. 2006;74:504–15.

23. Wise H, Hung C, Whiston E, Taylor J, Cole G. Extracellular ammonia at sites of pulmonary infection with Coccidioides posadasii contributes to severity of the respiratory disease. Microbial pathogenesis. 2013;59-60:19–28.

24. Reichard U, Hung C, Thomas P, Cole G. Disruption of the gene which encodes a serodiagnostic antigen and chitinase of the human fungal pathogen Coccidioides immitis. Infection and immunity. 2000;68:5830–8.

25. Xue J, Chen X, Selby D, Hung C, Yu J, Cole G. A genetically engineered live attenuated vaccine of Coccidioides posadasii protects BALB/c mice against coccidioidomycosis. Infection and immunity. 2009;77(8):3196–208.

26. Narra H, Shubitz L, Mandel M, Trinh H, Griffin K, Buntzman A, et al. A Coccidioides posadasii CPS1 Deletion Mutant Is Avirulent and Protects Mice from Lethal Infection. Infection and immunity. 2016;84:3007–16.

27. McGinnis M, Smith M, Hinson E. Use of the Coccidioides posadasii Deltachs5 strain for quality control in the ACCUPROBE culture identification test for Coccidioides immitis. Journal of clinical microbiology. 2006;44:4250–1.

28. Guevara-Olvera L, Hung C, Yu J, Cole G. Sequence, expression and functional analysis of the Coccidioides immitis ODC (ornithine decarboxylase) gene. Gene. 2000;242:437–48.

29. Mendoza BM, Saeger S, Campuzano A, Yu J, Hung C. Galleria mellonella Model of Coccidioidomycosis for Drug Susceptibility Tests and Virulence Factor Identification. Journal of fungi (Basel, Switzerland). 2024;10:ejof10020131.

30. Mead H, Hamm P, Shaffer I, Teixeira M, Wendel C, Wiederhold N, et al. Differential Thermotolerance Adaptation between Species of Coccidioides. Journal of fungi (Basel, Switzerland). 2020;6:ejof6040366.

31. Beyhan S, Gutierrez M, Voorhies M, Sil A. A temperature-responsive network links cell shape and virulence traits in a primary fungal pathogen. PLoS biology. 2013;11:e1001614.

32. Nguyen V, Sil A. Temperature-induced switch to the pathogenic yeast form of Histoplasma capsulatum requires Ryp1, a conserved transcriptional regulator. Proceedings of the National Academy of Sciences of the United States of America. 2008;105:4880–5.

33. Michielse C, van WR, Reijnen L, Manders E, Boas S, Olivain C, et al. The nuclear protein Sge1 of Fusarium oxysporum is required for parasitic growth. PLoS pathogens. 2009;5:e1000637.

34. Brown D, Busman M, Proctor R. Fusarium verticillioides SGE1 is required for full virulence and regulates expression of protein effector and secondary metabolite biosynthetic genes. Molecular plant-microbe interactions : MPMI. 2014;27:809–23.

35. Santhanam P, Thomma B. Verticillium dahliae Sge1 differentially regulates expression of candidate effector genes. Molecular plant-microbe interactions : MPMI. 2013;26:249–56.

36. Chen Y, Zhai S, Zhang H, Zuo R, Wang J, Guo M, et al. Shared and distinct functions of two Gti1/Pac2 family proteins in growth, morphogenesis and pathogenicity of Magnaporthe oryzae. Environmental microbiology. 2014;16:788–801.

37. Mirzadi GA, Mehrabi R, Robert O, Ince I, Boeren S, Schuster M, et al. Molecular characterization and functional analyses of ZtWor1, a transcriptional regulator of the fungal wheat pathogen Zymoseptoria tritici. Molecular plant pathology. 2014;15:394–405.

38. Okmen B, Collemare J, Griffiths S, der Burgt A van, Cox R, de WP. Functional analysis of the conserved transcriptional regulator CfWor1 in Cladosporium fulvum reveals diverse roles in the virulence of plant pathogenic fungi. Molecular microbiology. 2014;92:10–27.

39. Paes H, Derengowski L, Peconick L, Albuquerque P, Pappas GJ, Nicola A, et al. A Wor1-Like Transcription Factor Is Essential for Virulence of Cryptococcus neoformans. Frontiers in cellular and infection microbiology. 2018;8:369.

40. Tollot M, Assmann D, Becker C, Altmüller J, Dutheil J, Wegner C, et al. The WOPR Protein Ros1 Is a Master Regulator of Sporogenesis and Late Effector Gene Expression in the Maize Pathogen Ustilago maydis. PLoS pathogens. 2016;12:e1005697.

41. Vitter JS. Random Sampling with a Reservoir. ACM Trans Math Softw. 1985;11(1):37–57. doi:10.1145/3147.3165.

42. Engelthaler D, Roe C, Hepp C, Teixeira M, Driebe E, Schupp J, et al. Local Population Structure and Patterns of Western Hemisphere Dispersal for Coccidioides spp., the Fungal Cause of Valley Fever. mBio. 2016;7:e00550–16.

43. Teixeira M, Stajich J, Sahl J, Thompson G, Brem R, Dubin C, et al. A chromosomal-level reference genome of the widely utilized Coccidioides posadasii laboratory strain “Silveira”. G3 (Bethesda, Md). 2022;12:e6523976.

44. Smit A, Hubley R, Green P. RepeatMasker Open-4.0.; 2013-2015. Available from: http://www.repeatmasker.org.

45. Kirkland T, Muszewska A, Stajich J. Analysis of Transposable Elements in Coccidioides Species. Journal of fungi (Basel, Switzerland). 2018;4:ejof4010013.

46. Li H. Aligning sequence reads, clone sequences and assembly contigs with BWA-MEM; 2013.

47. Li H, Handsaker B, Wysoker A, Fennell T, Ruan J, Homer N, et al. The Sequence Alignment/Map format and SAMtools. Bioinformatics (Oxford, England). 2009;25:2078–9.

48. Walker B, Abeel T, Shea T, Priest M, Abouelliel A, Sakthikumar S, et al. Pilon: an integrated tool for comprehensive microbial variant detection and genome assembly improvement. PloS one;9:e112963.

49. Danecek P, Auton A, Abecasis G, Albers C, Banks E, DePristo M, et al. The variant call format and VCFtools. Bioinformatics (Oxford, England). 2011;27:2156–8.

50. Nguyen L, Schmidt H, von HA, Minh B. IQ-TREE: a fast and effective stochastic algorithm for estimating maximum-likelihood phylogenies. Molecular biology and evolution. 2015;32:268–74.

51. Kalyaanamoorthy S, Minh B, Wong T, von HA, Jermiin L. ModelFinder: fast model selection for accurate phylogenetic estimates. Nature methods. 2017;14:587–589.

52. Hoang D, Chernomor O, von HA, Minh B, Vinh L. UFBoot2: Improving the Ultrafast Bootstrap Approximation. Molecular biology and evolution. 2018;35:518–522.

53. Letunic I, Bork P. Interactive Tree Of Life (iTOL) v5: an online tool for phylogenetic tree display and annotation. Nucleic acids research. 2021;49:W293–W296.

54. Voorhies M, Cohen S, Shea T, Petrus S, Munoz J, Poplawski S, et al. Chromosome-Level Genome Assembly of a Human Fungal Pathogen Reveals Synteny among Geographically Distinct Species. mBio. 2022;13(1):e0257421.

55. R Core Team. R: A Language and Environment for Statistical Computing; 2022. Available from: https://www.R-project.org/.

56. Do C, Mahabhashyam M, Brudno M, Batzoglou S. ProbCons: Probabilistic consistency-based multiple sequence alignment. Genome research. 2005;15(2):330–40.

